# Chemogenetic tuning reveals optimal MAPK signaling for cell-fate programming

**DOI:** 10.1101/2024.12.19.629350

**Authors:** Brittany A. Lende-Dorn, Jane C. Atkinson, Yunbeen Bae, Kate E. Galloway

## Abstract

Cell states evolve through the combined activity of signaling pathways and gene regulatory networks. While transcription factors can direct cell fate, these factors rely on a cell state that is receptive to transitions in cell identity. How signaling levels contribute to the emergence of receptive cell states remains poorly defined in primary cells. Using a well-defined model of direct conversion, we examined how levels of the MAPK-activating oncogene HRAS^G12V^ influence direct conversion of primary fibroblasts to induced motor neurons. We demonstrate that an optimal ‘Goldilocks’ level of MAPK signaling efficiently drives cell-fate programming. Rates of direct conversion respond biphasically to increasing HRAS^G12V^ levels. While intermediate HRAS^G12V^ levels increase the rate of conversion, high levels of HRAS^G12V^ induce senescence. Through chemogenetic tuning, we set optimal MAPK activity for high rates of conversion in the absence of HRAS mutants. As MAPK pathways influence cell-fate transitions in development and disease, our results highlight the need to tune therapeutic interventions within a non-monotonic landscape that is shaped by genetics and levels of gene expression.

**Highlights:**

- MAPK signaling drives proliferation and conversion of fibroblasts to motor neurons
- Cell-fate programming responds biphasically to HRAS^G12V^ expression
- High HRAS^G12V^ expression induces senescence, which reduces conversion
- Chemogenetic tuning of MAPK activity increases conversion rates
- A small-molecule MAPK inducer drives high rates of conversion in the absence of HRAS^G12V^

## Introduction

The activity of the mitogen activated protein kinase (MAPK) pathway translates extracellular cues into changes in gene regulatory networks to shape cell-fate decisions.^1–6^ Pathway activity regulates critical cellular functions such as proliferation, differentiation, and survival. In the canonical MAPK cascade, the upstream RAS GTPase signals to RAF. Phosphorylation of RAF relays through the MAPK cascade from MEK to ERK, which translocates to the nucleus to regulate gene activity (Fig 1A). Nearly half of all tumors include activating mutations in receptor tyrosine kinases, RAS, or other MAPK pathway species.^7^ Among these MAPK mutations, RAS is the most frequently mutated, appearing in nearly a quarter of cancers.^8^ Mutations that lock RAS in its active GTP-bound state persistently stimulate downstream pathways to drive tumorigenesis.^9^ The strength and dynamics of MAPK signaling influence cell fate.^1,6^ Strong aberrant signaling triggers pathways that drive cells to terminal fates, whereas intermediate levels of signaling can support proliferation and self-renewal.^10–13^

**Figure 1.**
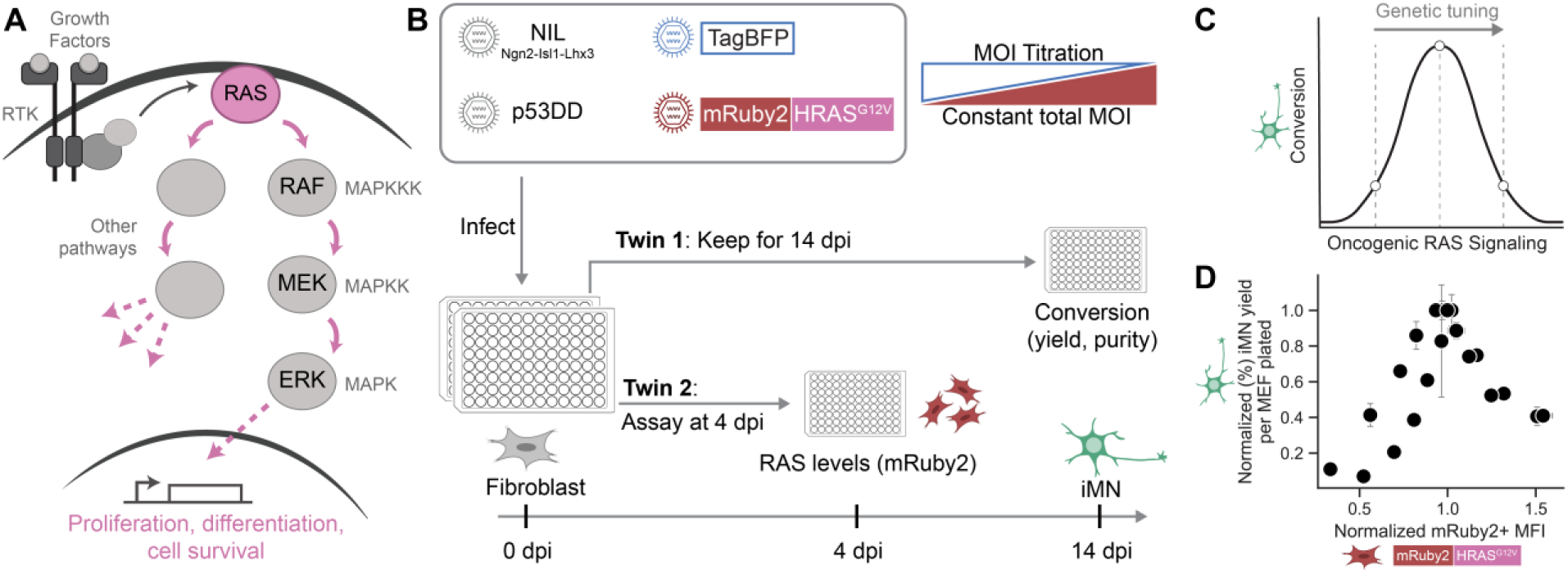
Cell-fate programming responds biphasically to titration of HRAS^G12V^. A. Simplified diagram of RAS signaling, with focus on the mitogen activated protein kinase (MAPK) pathway consisting of RAF, MEK, and ERK downstream of RAS. B. Schematic depicting viral delivery strategy of conversion cocktail. NIL (Ngn2, Isl1, and Lhx3 neuronal transcription factors) and p53DD (a p53 mutant) are delivered on retrovirus. mRuby2 tagged HRAS^G12V^ is delivered on lentivirus with expression driven by a CAG promoter. A “filler” lentivirus expressing TagBFP is included to keep lentivirus multiplicity of infection (MOI) constant for varying mRuby2-HRAS^G12V^ amounts. In a twin assay, two identical plates of cells are infected with the conversion cocktail. At 4 days post infection (dpi), mRuby2-HRAS^G12V^ expression levels are measured, and at 14 dpi the iMN yield and purity is quantified. C. Diagram depicting expected conversion results with mRuby2-HRAS^G12V^ MOI titration. D. Normalized iMN yield at 14 dpi vs. mRuby2-RAS+ geometric mean fluorescent intensity (MFI) at 4 dpi. Each point represents the mean of n = 3 technical replicates per bioreplicate ± standard error of mean (SEM). iMN yield is normalized so the maximum yield for each replicate overlays at 1.0 on the y-axis, and the mRuby2 levels are normalized so the mRuby2+ MFI at the MOI corresponding to the peak iMN yield overlays at 1.0 on the x-axis. n = 4 biological reps per condition.

While MAPK signaling contributes to diverse processes during cell-fate transitions, understanding how these pathways direct changes in cell identity remains challenging. Combinations of genetics and the dynamics of gene expression contribute to cell fate outcomes.^4,14–17^ Dynamic and non-monotonic relationships can obscure inference of causal relationships between profiles of gene expression and cell fates.^18–20^ In cellular reprogramming, addition of c-MYC, a downstream target of the MAPK pathway, increases rates of reprogramming to iPSCs.^21–23^ However, mild inhibition of MEK increases the total number of reprogrammed cells.^24^ In differentiation of embryonic stem cells (ESCs), attenuation of ERK signaling via MEK inhibition prohibits differentiation into neural lineages,^25,26^ whereas inhibition or knockdown of RSK1, a negative regulator of MAPK signaling, accelerates ESC differentiation.^27^ However, complete knockout of signaling nodes such as RAS and ERK1/2 in mouse ESCs results in growth arrest, limited differentiation, and apoptosis.^28,29^ *In vivo* and *in vitro*, the dynamics of MAPK signaling can dictate whether cells differentiate, die, or continue to divide.^1,6^ Notably, modulating ERK activity can rewire cell fates.^1^ Depending on the context, MAPK signaling can either increase or decrease rates of specific cell-fate transitions, suggesting complex processing of these dynamic signals.

Understanding the functional relationship between MAPK signaling and cell-fate transitions presents several challenges. Intrinsic and extrinsic variation can generate subpopulations that respond differently to cell-fate cues, obscuring connections between cell states and resulting cell fates.^19,30–33^ Even in genetically homogenous populations, the asynchronous and stochastic nature of cell-fate transitions makes observing and quantifying these events challenging.^34–36^ Consequently, many phenotypes that contribute to cell-fate transitions are studied in immortalized cell lines and stem cells, where genetic tools can be uniformly installed to map cell state to cell fate.^19,20,37–39^ However, the underlying genetic background for immortalization may limit the translatability of the findings from these systems to primary cells.

To investigate how healthy primary cells transition between somatic identities, we have recently developed system of direct conversion that generates induced motor neurons from fibroblasts at high rates.^30–32^ To improve our ability to track states that result in successful conversion, we improved the robustness and efficiency of conversion to generate larger numbers of converted cells.^30,31^ High rates of conversion offer the statistical power to draw inferences between cell states and cell fates in response to chemical and genetic perturbations.^30,31,40,41^ Further, we can accurately estimate the number of conversion events based simply on the number of neurons generated, which is not possible in transitions to mitotic cells. As post-mitotic cells, neurons do not divide. Thus, each neuron corresponds to exactly one conversion event. This direct conversion system provides a unique platform to systematically investigate the role of MAPK signaling across distinct stages of cell-fate transitions.

Inclusion of HRAS^G12V^ substantially increases rates of conversion of mouse embryonic fibroblasts.^30,31^ However, the specific impact of the levels of HRAS^G12V^ expression and the influence of MAPK signaling on direct conversion have not been previously defined. In this study, we systematically varied HRAS^G12V^ levels, explored the stoichiometric effects of multi-gene cassettes, and tested small-molecule regulators of the MAPK pathway to dissect their contributions to cell-fate programming. Our findings reveal a biphasic relationship between oncogenic RAS levels and conversion rates, demonstrating that high MAPK activity drives senescence, while moderate MAPK activity promotes proliferation and optimizes yield. By replacing HRAS^G12V^ with small-molecule inducer of MAPK signaling, we maintained high rates of conversion while eliminating oncogenic RAS. Overall, our results advance our understanding of the role of oncogenic RAS and MAPK signaling in our high efficiency direct conversion, offering insights for downstream translational applications.

## Results

### Cell-fate programming responds biphasically to titration of HRAS^G12V^

Oncogenes drive proliferation and cell-fate programming (Fig. 1A).^34,35,42–44^ To examine how the expression levels of MAPK mutants affect cell-fate transitions, we used a well-defined model of direct conversion.^31,32,41,45^ In this model system, transduction of mouse embryonic fibroblasts with a single cassette of transcription factors (Ngn2, Isl1, Lhx3 (NIL)) drives conversion to motor neurons (Fig. 1B).^31^ We can monitor conversion and quantify conversion events based on the activation of the motor neuron reporter, Hb9::GFP, from transgenic mouse embryonic fibroblasts. Unlike other cell-fate transitions, such as oncogenic transformation or conversion to mitotic cell types, direct conversion to a post-mitotic identity allows us to accurately quantify the number of conversion events based simply on the number of neurons.^31^

Introduction of transcription factors alone induces low rates of conversion (Fig. S1A-C). Expanding the population of hyperproliferative cells by introduction of mutant p53 (p53DD, a p53 mutant which lacks the DNA-binding domain), mutant RAS (HRAS^G12V^) and a small molecule TGFβ inhibitor (RepSox) dramatically increases conversion yield.^31,45^ This transient hyperproliferative population is highly receptive to transcription factors, converting at four-times the rates of other cells in the same conditions.^31,32^ By driving proliferation, levels of the mutant HRAS may influence the abundance and fate of this receptive population. To measure levels of the HRAS^G12V^, we added an N-terminal mRuby2 tag with a flexible GSG linker. To directly investigate how RAS expression levels affect conversion, we titrated the multiplicity of infection (MOI) of a lentivirus encoding mRuby2-HRAS^G12V^ (Fig. 1B, S2A-H). To ensure that conversion changes were not influenced by variations in total viral burden, we included a second lentivirus expressing TagBFP to maintain a constant viral load (i.e. total MOI) across conditions (Fig. 1B). As expected, the level of mRuby2-HRAS^G12V^ increases with MOI (Fig. S2C).

Previous work indicates that proliferation of epithelial cells increases non-monotonically in response to MAPK signaling.^4^ Thus, we hypothesized that conversion would respond non-monotonically to increasing levels of mRuby2-HRAS^G12V^ (Fig. 1C). To define how levels of HRAS^G12V^ affect cell-fate programming, we used a twin-plate assay to map expression levels of HRAS^G12V^ to conversion events. In a twin-plate assay, we measure mRuby2-HRAS^G12V^ levels at 4 days post-infection (dpi) from one plate of cells and measure conversion via Hb9::GFP activation at 14 dpi from a second “twin” plate (Fig. 1B).

Rapidly proliferating cells are smaller and show lower levels of transgene expression.^31,32^ To control for proliferation-mediated differences, we normalized the mRuby2 fluorescence intensities and conversion yields across conditions and replicates (Fig. S2D-H). Normalizing by replicate allows us to control for the combined batch effects of virus and primary mouse embryonic fibroblasts. We normalized each replicate to the MOI corresponding to the peak iMN yield within that replicate. This normalization reveals a clear biphasic relationship between mRuby2-HRAS^G12V^ levels and conversion yield (Fig. 1D). These results indicate that optimal conversion rates are driven by intermediate levels of RAS expression. Additionally, we examined the effect of mRuby2-HRAS^G12V^ levels on conversion with a SNAP-tagged p53DD that allows us to visualize p53DD expression in the presence of a fluorescently labeled SNAP substrate. While both variants show a biphasic conversion response, SNAP-DD increases the total yield of neurons and reduces the absolute optimal levels of mRuby2-HRAS^G12V^ (Fig. S1A-C, Fig S1A-D). Thus, the exact optimal level of RAS expression depends on the properties of the p53 mutant (Fig. 1D, S1D, S2E-F). Together our data indicates that an optimal level of HRAS^G12V^ drives cell-fate transitions. The identity and properties of genetic variants can tune the optimum HRAS^G12V^ level and the rates of conversion.

### HRAS^G12V^ produces biphasic conversion through MAPK signaling and proliferation

Overexpression of HRAS^G12V^ drives MAPK signaling.^8,46^ Thus, we hypothesized that increasing HRAS^G12V^ expression increases rates of proliferation rates and conversion by inducing MAPK signaling (Fig. 2A). To limit extrinsic variation associated with different copy numbers, we encoded both HRAS^G12V^ and p53 mutants on a single viral transcript upstream and downstream of an Internal Ribosomal Entry Site (IRES). The relative ordering of genes around an IRES can generate different ratios of transgene expression while controlling for relative copy number.^47^ Therefore, we expected to observe a range of HRAS^G12V^ levels and conversions rates across cassettes with minimal genetic sources of extrinsic variance.

**Figure 2.**
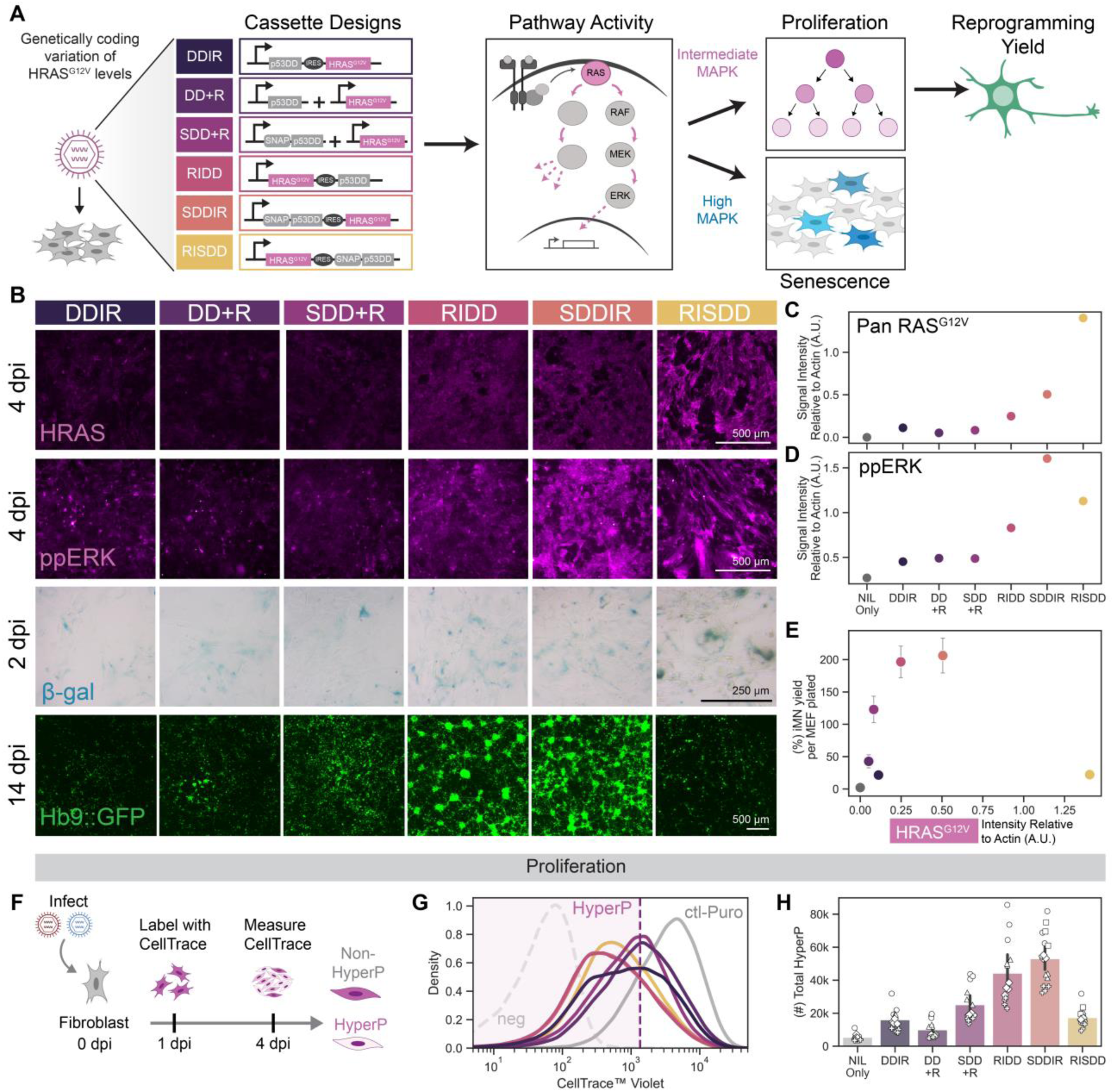
HRAS^G12V^ produces biphasic conversion through MAPK signaling and proliferation. A. Schematic depicting polycistronic cassette designs expressing HRAS^G12V^ and either p53DD or SNAP-p53DD, separated by an internal ribosomal entry site (IRES). Transduced cells were analyzed for phenotypes that are expected to change with varying RAS expression levels. Conversion conditions always include NIL transcription factors. B. Representative images of immunofluorescent staining for HRAS and ppERK at 4 days post infection (dpi), cells stained for senescence-associated β-galactosidase (β-gal) at 2 dpi, and Hb9::GFP expression in iMNs at 14 dpi. Scale bar represents 500 μm for HRAS, ppERK, and Hb9::GFP images, and 250 μm for β-gal images. C. Pan RAS^G12V^ expression levels quantified from a western blot and normalized to β-Actin levels. D. ppERK levels quantified from a western blot and normalized to β-Actin levels. E. iMN yield at 14 dpi vs. Pan RAS^G12V^ expression normalized to β-Actin levels measured from a western blot. Mean iMN yield is shown ± standard error of mean (SEM). F. Diagram depicting cell proliferation assay. Cells are labeled with CellTrace dye at 1 dpi, and fluorescent signal is measured at 4 dpi. Cells with a lower CellTrace signal have proliferated more than cells with a higher signal. G. Representative histograms of CellTrace levels at 4 dpi across conditions. Hyperproliferative (HyperP) cells are gated at the 20% lowest CellTrace signal in a control Puro (ctl-Puro) infected condition for a given biological replicate. H. Total number HyperP cells at 4 dpi across conditions. Mean is shown with 95% confidence interval; marker style denotes biological reps; n = 4 biological reps per condition.

We compared the four different single-transcript designs to the dual-virus delivery conditions (Fig 2A). To measure HRAS^G12V^ expression levels across all conditions, we used immunofluorescent staining (Fig. 2B) and western blot quantification (Fig. 2C, S3A-E). At 4 dpi, we observed a wide range of HRAS^G12V^ levels across conditions. By western blot, overexpression of HRAS^G12V^ ranges from 2 to 20 times the endogenous Pan RAS levels (Fig. S3A, C). When we order conditions by levels of HRAS^G12V^ expression, we find a biphasic relationship between HRAS^G12V^ levels and conversion (Fig. 2B, S3F-G). MAPK signaling increases the phosphorylation of the downstream MAP kinase, ERK1/2. To examine MAPK signaling activity across conditions, we measured levels of phosphorylated ERK1/2 (ppERK). As expected, both western blot and immunofluorescence imaging indicates that levels of ppERK correlate with levels of HRAS^G12V^ (Fig. 2B-D). At the highest levels of HRAS^G12V^, we observe substantial increases in β-actin staining, reducing the normalized signal of ppERK (Fig. 2D, Fig. S3B). Increases in β-actin align with changes we observe in cell size at high levels of HRAS^G12V^and reports of proteome remodeling in large cells.^48^ Plotting the HRAS^G12V^ levels—as quantified from a western blot—against the iMN yield at 14 dpi confirmed the biphasic response (Fig. 2E, S3H), consistent with the results from the mRuby2-HRAS^G12V^ MOI titration (Fig. 1D). Together these data indicate that optimal levels of HRAS^G12V^ induce an optimal level of MAPK signaling for high rates of conversion.

Proliferation drives cells to a state that is highly receptive to lineage-specifying transcription factors.^31,32,45^ As MAPK signaling induces cellular proliferation, we hypothesized that optimal levels of HRAS^G12V^ expression support conversion by increasing the population of hyperproliferative cells. To assess proliferation history across conditions, we performed a CellTrace dye dilution assay.^31,32^ Cells were labeled at 1 dpi, and CellTrace signal was analyzed at 4 dpi using flow cytometry (Fig. 2F). Lower CellTrace signal indicates a history of more proliferation. We denote hyperproliferative (HyperP) cells as the 20% lowest CellTrace signal in the control condition (Fig. 2G). As expected from the conversion rates, we observe that increasing HRAS^G12V^ levels increases the total number of HyperP cells up to a point. Above this threshold, increasing HRAS^G12V^ reduces the number of HyperP cells and the conversion rate (Fig. 2H, S3F, H-I). In particular, we found that the RISDD condition, which had the highest HRAS^G12V^ and elevated MAPK signaling levels, consistently produces lower rates of conversion.

In diverse primary cells, overexpression of mutant HRAS leads to oncogene-induced senescence.^49–57^. Senescence is a state of permanent cell cycle arrest associated with metabolic changes.^58^ Staining for β-galactosidase (β-gal), a senescence marker, reveals an increase in cells with a strong β-gal signal in the RISDD condition (Fig. 2B). Morphologically, we observed that cells with strong β-gal signal are very large and flat (Fig. S3J), another signature of senescence.^59^ We conclude that the high HRAS^G12V^ levels in the RISDD condition induce senescence, preventing the cell cycle progression necessary for conversion.

Our results demonstrate that low levels of HRAS^G12V^ limit MAPK signaling, proliferation and conversion. Conversely, excessive HRAS^G12V^ expression elevates MAPK signaling to levels that induce senescence. This balance between proliferation and senescence creates a biphasic response of cell fate to HRAS^G12V^ levels.

### Tuning MAPK signaling attenuates senescence and increases conversion

Putatively, high levels of HRAS^G12V^ drive oncogene-induced senescence through MAPK signaling. At the highest levels of oncogenic RAS expression, inhibition of MAPK signaling may rescue conversion (Fig. 3A). To test this hypothesis, we treated cells expressing high levels of HRAS^G12V^ with PD0325901, a MEK inhibitor (MEKi), from 1-14 dpi. MEKi blocks MAPK signaling downstream of RAS by inhibiting MEK1/2-mediated transmission of signaling to ERK1/2.

**Figure 3.**
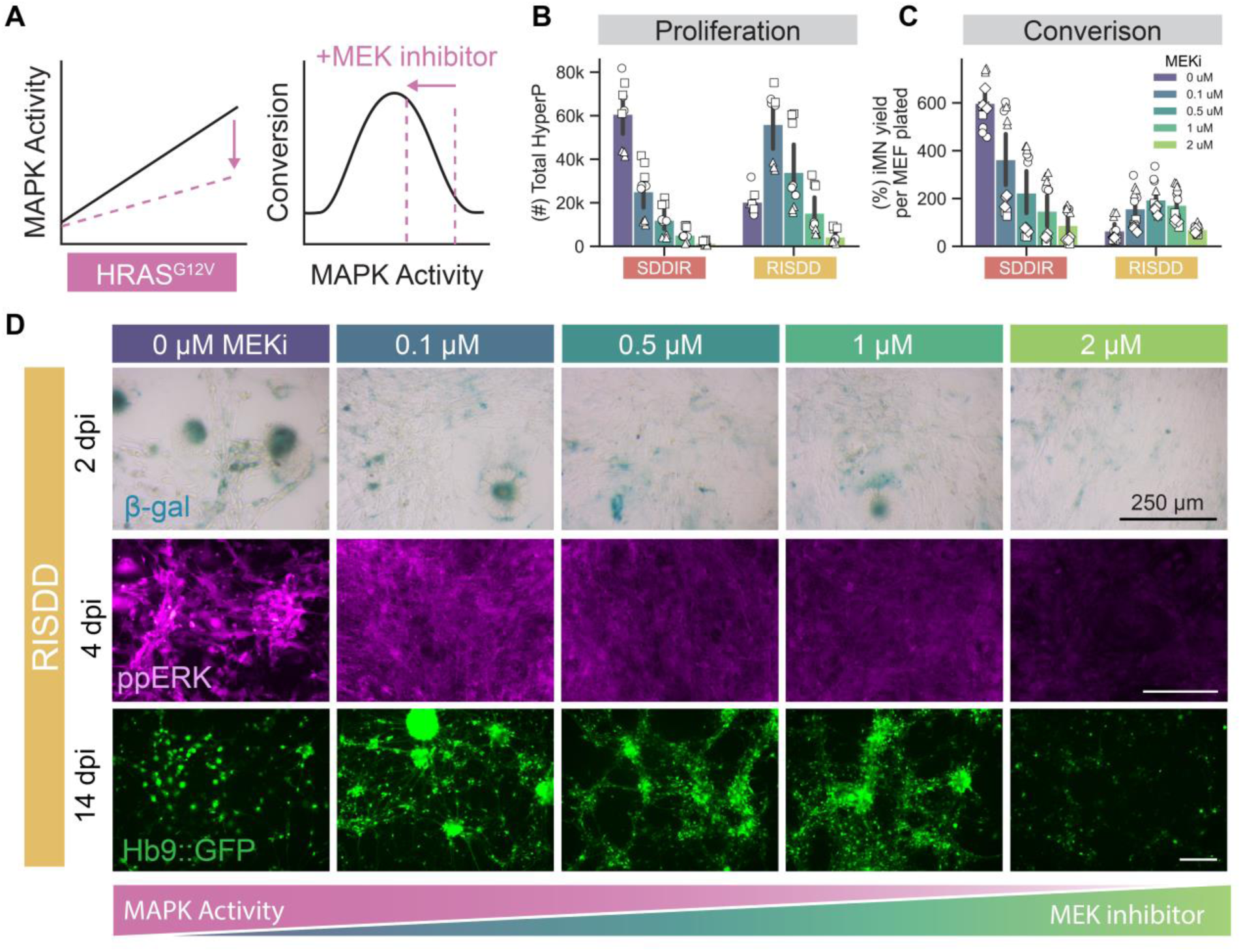
Tuning MAPK signaling attenuates senescence and increases conversion. A. Diagram depicting expected results of adding a MEK inhibitor on MAPK signaling levels and conversion. B. Total number HyperP at 4 dpi with a MEK inhibitor (PD0325901) titration for the two polycistronic cassettes with highest HRAS^G12V^ expression (SDDIR and RISDD). Mean is shown with 95% confidence interval; marker style denotes biological reps; n = 3 biological reps per condition. C. Conversion yield at 14 dpi with a MEK inhibitor titration for the two polycistronic cassettes with highest HRAS^G12V^ expression. Mean is shown with 95% confidence interval; marker style denotes biological reps; n = 4 biological reps per condition. D. Representative images of cells stained for senescence-associated β -galactosidase (β-gal) at 2 dpi, ppERK at 4 dpi, and Hb9::GFP expression in iMNs at 14 dpi for the polycistronic cassette with the highest HRAS^G12V^ expression (RISDD) with MEKi titration. Scale bar represents 250 μm.

We compared the effect of MEKi at two levels of HRAS^G12V^ expression; intermediate and high. At high levels of HRAS^G12V^ expression, we expect that MEKi will increase proliferation and conversion. From the model of conversion following a biphasic response, we expect that MEKi treatment at intermediate levels of HRAS^G12V^ will reduce conversion (Fig. 3A). In conversion, we used two polycistronic cassettes that code for identical proteins but generate intermediate and high levels of HRAS^G12V^ expression, SDDIR and RISDD, respectively (Fig. 3B-D). We confirmed that MEKi reduces ppERK levels at 4 dpi in a dose-dependent manner (Fig. 3D, S4A-D). At intermediate levels of HRAS^G12V^ (i.e. the SDDIR condition), MEKi treatment reduces the number and fraction of HyperP cells at 4 dpi and the rate of conversion (Fig. 3B-D, S4E-F). Conversely, at the highest levels of HRAS^G12V^ (i.e. the RISDD condition), low concentrations of MEKi increase proliferation and conversion. Thus, MEKi treatment rescues conversion yield at high HRAS^G12V^ expression while MEKi reduces conversion yield at intermediate levels of HRAS^G12V^ expression. Together these data reinforce the idea that optimal levels of HRAS^G12V^expression drive conversion by tuning MAPK activity to the optimal regime.

Inhibiting MAPK activity at high levels of HRAS^G12V^ expression may promote conversion by driving proliferation while limiting senescence. To investigate whether MEKi mitigates HRAS^G12V^-induced senescence, we treated cells expressing high levels of HRAS^G12V^ (i.e. RISDD condition) with varying MEKi concentrations at 1 dpi. We stained for senescence-associated β-gal at 2 dpi. MEKi reduces the number of cells showing strong β-gal signal (Fig. 3D). At 14 dpi, iMNs do not show strong β-gal signal (Fig. S4G), indicating that β-gal staining is unrelated to the iMN state.

Recognizing that RAS activates other signaling pathways, we tested whether inhibiting the PI3K or JNK pathways affects the rate of conversion in the RISDD condition. Immunofluorescent staining for phospho-AKT (pAKT) in the polycistronic cassettes (Fig. 2A) reveals that pAKT levels increase with increasing HRAS^G12V^ levels (Fig. S5A). Therefore, we treated cells with an AKT inhibitor (AKTi) to block PI3K signaling at high levels of HRAS^G12V^ (Fig. S5B). At low AKTi doses, we found no improvement in proliferation or conversion; higher doses reduce cell counts, suggesting toxicity (Fig. S5C-E). AKTi treatment fails to attenuate senescence (Fig. S5E). Depending on the cellular context, JNK signaling can prevent or induce senescence.^60^ We tested a JNK inhibitor (JNKi). In the RISDD condition, JNKi treatment does not attenuate senescence (Fig. S5F).

When HRAS^G12V^ expression is excessively high in the RISDD condition, inhibiting the MAPK pathway with MEKi effectively rescues cell fate programming. Tuning MAPK signaling increases proliferation and reduces senescence. In contrast, inhibition of the PI3K or JNK pathways does not produce similar effects. Thus, driven by high HRAS^G12V^ expression, high levels of MAPK signaling induces senescence, which we can tune by MEKi treatment.

### Activation of MAPK signaling induces high rates of conversion in the absence of mutant RAS

We wondered if activation of MAPK signaling — in the absence of a RAS mutant — could drive high efficiency conversion. Diverse extracellular cues, including small molecules, stimulate MAPK signaling.^61–63^ Potentially, transient stimuli such as small molecules could replace delivery of the oncogene HRAS^G12V^, increasing safety and simplifying the genetic cocktail for direct conversion. The small molecule phorbol 12-myristate 13-acetate (PMA) activates protein kinase C to trigger signaling in the MAPK pathway.^1,64–66^ We hypothesized that addition of PMA to converting cells would drive direct conversion by inducing signaling and proliferation, offering an alternative to activation via oncogenic RAS expression (Fig. 4A-B). Across a range of concentrations, we find that addition of PMA increases the levels of ppERK (Fig. 4C, S6A-D).

**Figure 4.**
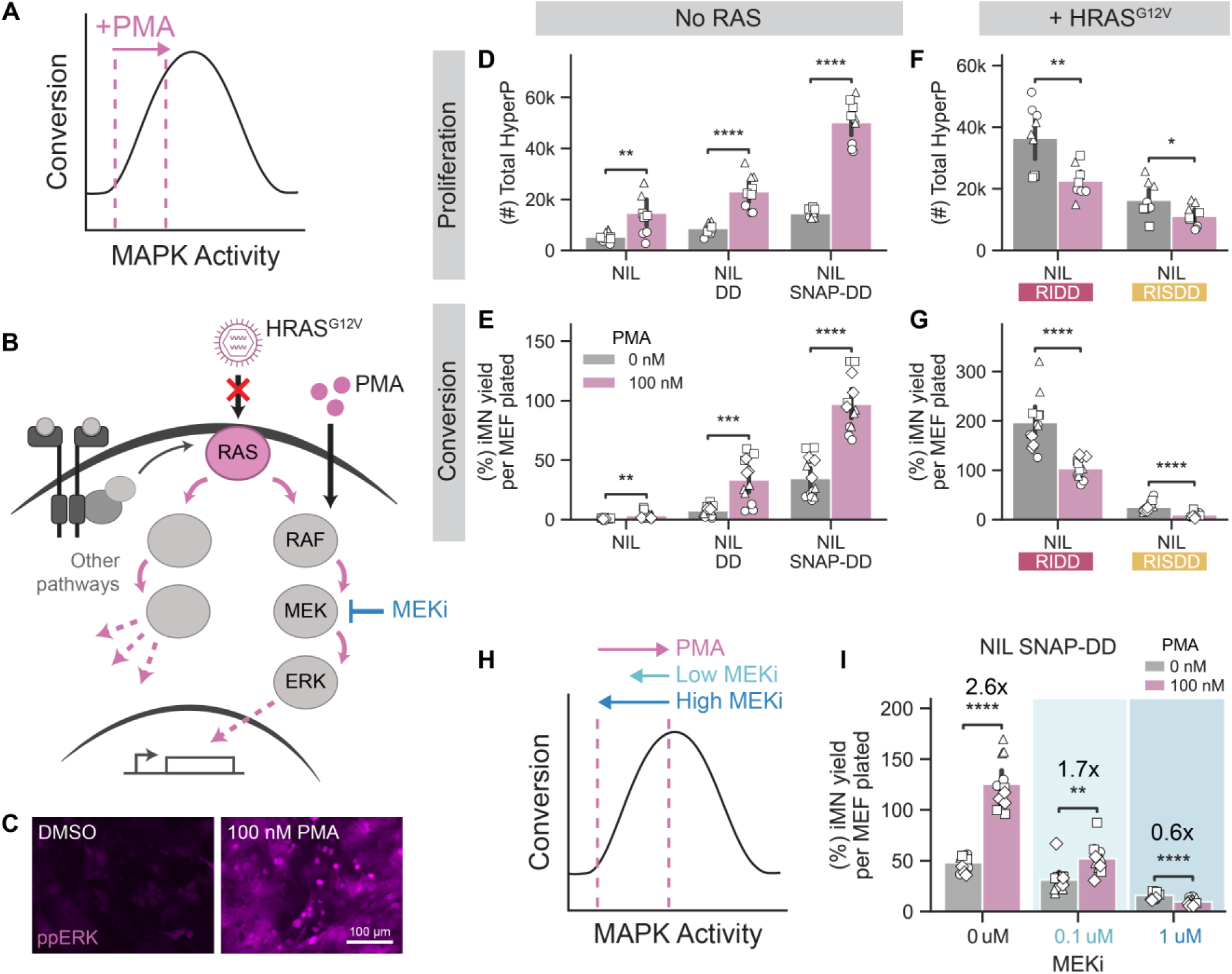
Activation of MAPK signaling induces high rates of conversion in the absence of mutant RAS. A. Diagram depicting expected results of adding PMA, a small molecule activator of the MAPK pathway, on MAPK signaling levels and conversion. B. Schematic depicting the replacement of viral transduction of RAS oncogene with small molecule PMA to activate MAPK signaling. C. Representative images of immunofluorescent staining for ppERK in cells fixed 20 minutes after treatment with either a vehicle control (DMSO) or 100 nM PMA. Scale bar represents 100 μm. D-E. Total number HyperP cells at 4 dpi (D) and conversion yield at 14 dpi (E) with 100 nM PMA treatment for conditions without RAS. Mean is shown with 95% confidence interval; marker style denotes biological reps; n = 3 or 4 biological reps per condition; t-test independent samples. F-G. Total number HyperP cells at 4 dpi (F) and conversion yield at 14 dpi (G) with 100 nM PMA treatment for conditions with polycistronic cassettes containing HRAS^G12V^. Mean is shown with 95% confidence interval; marker style denotes biological reps; n =3 or 4 biological reps per condition; t-test independent samples. H. Diagram depicting expected results of adding PMA in combination with a MEK inhibitor to test whether the effects of PMA are specific to the MAPK pathway. I. Conversion yield quantified at 14 dpi for cells infected with NIL SNAP-DD and treated with combinations of 0 nM or 100 nM PMA to activate MAPK signaling and 0 μM, 0.1 μM, or 1 μM of MEK inhibitor (PD0325901) to inhibit MAPK signaling. Small molecules were added to the media beginning at 1 dpi. Mean is shown with 95% confidence interval; marker style denotes biological reps; n = 4 biological reps per condition; t-test independent samples. Significance summary: p > 0.05 (ns); *p ≤ 0.05; **p ≤ 0.01; ***p ≤ 0.001; and ****p ≤ 0.0001.

To explore the effects of PMA on proliferation and conversion, we treated converting cells with PMA from 1 dpi through 14 dpi. In the absence of mutant HRAS, PMA treatment increases both the number and percentage of HyperP cells at 4 dpi (Fig. 4D, S6E-F). The boost in proliferation correlates with improved iMN yield and purity (Fig. 4E, S6G-H). Specifically, PMA increases conversion yield by up to 5-fold. Compared to transcription factors alone, addition of 100 nM PMA with SNAP-DD increases conversion yield by 120-fold, approaching 100% yield. However, PMA-converted iMNs have less mature morphology and neurite outgrowth compared to HRAS^G12V^-converted cells (Fig. S6D). Higher concentrations of PMA do not increase yield and at the highest concentration, we observed lower conversion rates (Fig. S6G-H).

If PMA and HRAS^G12V^ work through a shared mechanism of optimal MAPK signaling, we would expect that addition of PMA to conditions with high HRAS^G12V^ would not increase the rates of proliferation or conversion. When we added PMA to converting cells with high levels of HRAS^G12V^, we observed no increase in proliferation (Fig. 4F, S6I-J) or conversion (Fig. 4G, S6K-L). Rather, both metrics show slight reductions. As PMA does not increase rates of conversion in HRAS^G12V^-expressing conditions, our results suggest that both PMA and HRAS^G12V^ drive conversion through MAPK signaling.

To demonstrate that PMA increases rates of conversion by inducing MAPK signaling, we treated cells infected with NIL SNAP-DD with combinations of PMA and MEKi at different concentrations (Fig. 4H). Low MEKi concentration partially reduces the PMA-mediated boost in conversion, while high concentration of MEKi completely abolishes the boost in conversion (Fig. 4I, S6M). Overall, we demonstrate that we can replace oncogenic RAS with a small molecule activator of MAPK signaling to induce high rates of conversion.

## Discussion

Understanding how signaling pathways influence cell-fate transitions in healthy primary cells offers a window into development, tissue homeostasis, and disease. In this work, we demonstrate that cell-fate programming of primary cells responds biphasically to MAPK signaling. Using a combination of chemical and genetic activators to tune signaling, we find that intermediate levels of signaling promote proliferation and conversion of primary mouse embryonic fibroblasts to motor neurons without inducing senescence. Co-expressed MAPK and p53 mutants tune MAPK signaling to set the level optimal for proliferation and conversion. Rates of conversion vary as a function of genetic factors, such as differences in p53 status,^51,67,68^ and extrinsic features such as the batch of virus or cells. Importantly, when we applied chemical activators and inhibitors to the MAPK pathway, we observe diverging responses that are dependent on the expression level of mutant HRAS. Understanding this non-monotonic landscape allowed us maintain high rates of conversion in the absence of HRAS^G12V^ by supplementing with a small-molecule activator of the MAPK pathway. Extending these finding to additional systems and human cells may reveal opportunities to precisely reshape signaling and selectively control cell-fate transitions, improving the precision of therapeutic interventions.

Our findings align with other studies that show that an optimal level of MAPK signaling promotes transformation and tumorigenesis.^69–71^ Directly changing levels of expression of MAPK pathway mutants reveals that intermediate levels of mutant expression support proliferation.^4,52^ Interestingly, we observe that inhibition of MAPK signaling promotes survival in cells with high expression of HRAS^G12V^. Attenuated signaling in the presence of high levels of HRAS^G12V^ limits oncogene-induced senescence. High MAPK signaling may generally drive cells to terminal fates. Strong MAPK pathway-activating mutations, such as in KRAS and BRAF, are mutually exclusive in cancer.^10^ Simultaneous expression of these mutants is cytotoxic to cells and suppresses tumor formation.^11,12^ In primary mouse embryonic fibroblasts, co-expression of hyperactive KRAS and BRAF mutants halts proliferation and causes oncogene-induced senescence.^11^ Similar to our findings, inhibition of MAPK signaling can reduce senescence and toxicity associated with oncogene overexpression.^5,12^ Conversely, MAPK pathway activation may reduce proliferation and increase cell stress, leading to senescence or cell death, presenting a potential treatment strategy for cancers with hyperactive MAPK signaling.^72–74^ Resistance to MAPK pathway inhibitors could be mitigated by removing the inhibitor, resulting in toxic hyperactivation of MAPK signaling.^75,76^ Our findings provide a nuanced perspective on MAPK modulation that may better guide development of tailored MAPK inhibitors for therapeutic contexts. Tools that precisely control the expression of oncogenes may support the development of genetic systems that more accurately model cell states prior to and during transformation.^77,78^

Addition of the small-molecule MAPK activator PMA allowed us to generate neurons at high rates in the absence of mutant HRAS. Elimination of oncogenes reduces the risk of transformation, improving the safety profile of scalably-produced, *in vitro*-derived cells for translational applications such as cellular therapies. While PMA improves conversion efficiency without oncogenic RAS, the PMA-converted cells exhibit less mature morphologies compared to their RAS-converted counterparts (Fig. S6D). Constant stimulation of MAPK signaling may impede neurite outgrowth. In development and differentiation, levels and dynamics of MAPK signaling impact neural differentiation and neurite outgrowth.^1,79–81^ By tuning the duration of PMA stimulation, we may be able to increase neuronal morphology and maturation while retaining high rates of conversion. Alternatively, PMA-converted cells may require a longer conversion timeline to mature, which we can explore through prolonged culture in the presence or absence of PMA. While the safety profile of the PMA-based protocol is an improvement over oncogenic RAS, more work is needed to assess the long-term stability and translational potential of PMA-converted cells.

Both HRAS^G12V^ and PMA improve conversion through MAPK pathway activation, yet their distinct effects suggest opportunities to explore additional MAPK modulators to increase cell yields and morphological maturity. While HRAS^G12V^ effectively drives conversion of MEFs to iMNs, HRAS^G12V^ fails to support conversion of adult human fibroblasts to neurons.^31^ Addition of myr-AKT, BCL2, and c-MYC with NIL and p53DD in primary adult human fibroblasts increases rates of conversion.^31^ Notably, AKT and MYC are both activated downstream of RAS signaling, suggesting that species-specific signaling patterns may differentiate the effects of mutant RAS on mouse and human cells. RAS isoforms differ in their mutation frequency and endogenous expression levels across human cancers. KRAS is the most frequently mutated isoform, followed by NRAS and HRAS.^8,82^ Other RAS mutants may have different optimal expression levels based on their transforming potential and cell context.^46^ Differences in the response of adult human fibroblasts and mouse embryonic fibroblasts to HRAS^G12V^ may reflect a distinct window of sensitivity to RAS levels, species-specific disparities, or differences in developmental stage of the initial cells. Investigating these differences could shed light on the molecular requirements for conversion of human cells and pave the way for improved protocols.

In summary, HRAS^G12V^ can drive cells to a state receptive to conversion by promoting proliferation through MAPK signaling. However, extremely high expression levels of mutant RAS lead to oncogene-induced senescence. This tradeoff between proliferation and senescence results in a biphasic correlation between HRAS^G12V^ expression and conversion. We can direct cells through MAPK signaling regimes by genetically tuning HRAS^G12V^ levels or via small-molecule interventions. Overall, our work underscores the critical role of fine-tuning MAPK signaling during cell-fate transitions.

## Author Contributions

B.A.L and K.E.G. conceived and outlined the project. B.A.L. cloned DD RAS polycistronic cassette retrovirus plasmids and mRuby2-RAS and TagBFP lentivirus plasmids. B.A.L., J.C.A., and Y.B. performed reprogramming experiments and analyzed data. B.A.L. and J.C.A. performed immunofluorescent staining. B.A.L. performed western blots and senescence staining. B.A.L. and K.E.G. wrote the manuscript. K.E.G. supervised the project.

## Acknowledgements

Research reported in this manuscript was supported by the National Institute of General Medical Sciences of the National Institutes of Health under Health under award number R35-GM143033, by the National Science Foundation under the NSF-CAREER under award number 2339986, and the with funding from Institute for Collaborative Biotechnologies. B.A.L. is supported by the National Science Foundation Graduate Research Fellowship Program under grant No. 1745302. We thank Emma Peterman, Sneha Kabaria, Kasey Love, and Adam Beitz for feedback on the development of the manuscript.

## Data and Materials Availability

Raw data and python data analysis code is available from the corresponding author upon request.

## Materials and Methods

### Cell lines and tissue culture

Mouse embryonic fibroblasts (MEFs), Platinum-E (Plat-E), and HEK293T cells were cultured in Dulbecco’s Modified Eagle Medium (DMEM) (Genesee Scientific, 25-501) supplemented with 10% fetal bovine serum (FBS) (Genesee Scientific, 25-514H) and incubated at 37°C and 5% CO_2_. Every 3 passages, Plat-Es were selected with 10 µg/mL blasticidin and 1 µg/mL puromycin. Cells were counted using a hemocytometer when seeding. All cells were routinely tested for mycoplasma contamination.

### MEF dissection and isolation

E14.5 embryos were harvested from mice after crossing a heterozygous Hb9::GFP reporter mouse with a wild type (C57BL/6) mouse. Hb9::GFP-positive embryos were identified using a blue laser to detect GFP expression in the spinal cord. The head and internal organs were removed from the embryo, followed by mechanical dissociation of the remaining tissue with razor blades. The tissue was further dissociated using 0.25% trypsin-EDTA (Genesee Scientific, 25-510), first using razor blades and then via trituration. After neutralization with DMEM + 10% FBS, centrifugation, and resuspension, the cells were passed through a 40 μm cell strainer to obtain a single cell suspension. Cells were plated onto 0.1% gelatin coated 10 cm dish (1 embryo per dish). At ∼80% confluency, each dish was dissociated split onto 3 gelatin coated dishes. Once the dishes reached ∼80% confluency again, the MEFs were cryopreserved in 90% FBS and 10% DMSO and stored in liquid nitrogen. MEFs were tested for mycoplasma before use.

### Plasmid construction

Plasmids were constructed using Gibson and Golden Gate cloning methods. Retroviral plasmids were cloned via LR recombination into the pMXs-WPRE-DEST plasmid. Lentiviral plasmids were cloned via LR into a pLentiX1-Harbor plasmid. All viral plasmids were confirmed using whole plasmid sequencing or Sanger sequencing.

### Retroviral transduction and conversion of MEFs to iMNs

For retrovirus production, Plat-Es were seeded at 0.9 million per well onto 6-well plates coated with 0.1% gelatin for at least 5 minutes. The next day, Plat-Es were transfected with 1.8 μg of transfer plasmid per 6-well using a 4:1 ratio of µg PEI:µg DNA in KnockOut™ DMEM (ThermoFisher Scientific, 10-829-018). After 18 hours, the media was replaced with 1.25 mL of 25 mM HEPES buffered DMEM + 10% FBS. 24 and 48 hours later, the viral supernatant was collected and filtered using 0.45 μm PES syringe filters. Culture media was replenished after the first virus collection.

Primary MEFs were thawed from cryopreserved stocks three days prior to transduction into a gelatin coated T-75 flask, and seeded at 10k cells per 96-well onto gelatin coated plates at 1 day prior to transduction. The next two days, MEFs are transduced with freshly collected and filtered virus. Each 96-well receives 11 μL of each virus for a given condition diluted in media to a final volume of 100 μL per 96-well, with a final concentration of 5 μg/mL polybrene (Sigma-Aldrich, H9268) to increase transduction efficiency. At 1 day post infection (1 dpi), virus-containing media is replaced with fresh DMEM + 10% FBS.

At 3 dpi, the culture media was replaced with N3 media (DMEM/F12 (Fisher Scientific, 21331020) containing N2 (Fisher Scientific, 17-502-048), B27 (Thermo Scientific, 17504044), and 1% Glutamax (Thermo Fisher Scientific, 35050061); neurotropic growth factors – BDNF, GDNF, CNTF, and FGF (R&D Systems) – were added immediately before use to a final concentration of 10 ng/mL). Experimental conditions with HRAS^G12V^ also included the small molecule RepSox (Selleck Chemicals, S7223) at 7.5 μM starting at 3 dpi. N3 media is replaced every 2 to 3 days until 14 dpi. At 14 dpi, cells were dissociated using DNase and papain. One vial each of DNase and papain (Worthington Biochemical, LK003172 and LK003178) were reconstituted in 8 mL of DMEM/F12. 40 μL of DNase/papain solution was used per 96-well and allowed to incubate at 37°C for ∼15 minutes or until cells detached. After dissociation, cells were resuspended in 250 μL PBS for analysis via flow cytometry. iMNs were gated on a population of bright Hb9::GFP+ cells. Dim Hb9::GFP+ cells were excluded from iMN quantification. iMN yield and purity were calculated as follows:

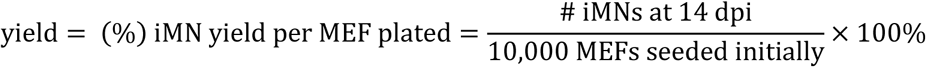

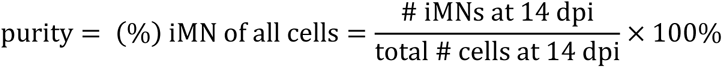

### Lentiviral production in HEK293Ts

HEK293T cells were seeded at 6 million per 10 cm dish coated with 0.1% gelatin. The next day, each plate of 293Ts were co-transfected with 6 µg packaging plasmid (psPax2, Addgene #12260), 12 µg envelope plasmid (pMD2.G / VSVG, Addgene #12259), and 6 µg of lentivirus transfer plasmid using a 4:1 ratio of μg PEI:μg DNA. 6-8 hours later, the media was replaced with 6.5 mL of 25 mM HEPES buffered DMEM + 10% FBS. 24 and 48 hours after the media change, viral supernatant was collected and stored at 4°C, replenishing with fresh media after the first collection. After the second collection, viral supernatant was filtered with 0.45 μm PES filters, mixed with Lenti-X concentrator (Takara Bio, 631232) at a 3:1 volume ratio of viral supernatant:Lenti-X, and then stored overnight at 4°C. The next day, the virus mixtures were centrifuged at 1500 × g for 45 minutes at 4°C to pellet the virus. After removing the supernatant, the pellet was resuspended in 250 μL of media per original 10 cm plate. Concentrated virus was either kept at 4°C for less than a week, or stored at −80°C for longer periods.

### Functional titer measurement and quantification

MEFs were seeded at 10k per well onto 96-well plates coated with 0.1% gelatin for at least 5 minutes. Two days later, cells were transduced with a serial dilution of lentivirus. The starting volume of virus was 5 μL of concentrated lentivirus per 96-well, with a 4x series dilution into 4 more wells for a total of 5 virus concentrations. Lentivirus was diluted in DMEM + 10% FBS so that each well had a final volume of 100 μL per 96-well, with a final concentration of 5 μg/mL polybrene to increase transduction efficiency. After adding the virus, the cells were spinfected by centrifuging the plate at 1500 × g for 30 minutes at 32°C to further increase transduction efficiency. 24 hours later at 1 dpi, the virus containing media was replaced with fresh DMEM + 10% FBS. At 2 dpi, the cells were dissociated using trypsin, and the infection efficiency was measured via flow cytometry.

The fraction of fluorescent-positive cells was calculated for each well. To ensure single-integration events, only wells with infection rates below 40% were used for titer calculations. The functional titer in transducing units (TU) per μL was then calculated as follows:

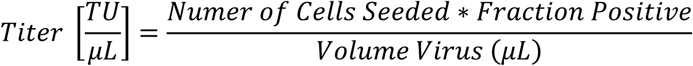

Functional titer values were used to calculate the volume of concentrated lentivirus needed for specific multiplicities of infection (MOIs). For example, seeding 10k MEFs in a 96-well required 10,000 TU for MOI 1 or 50,000 TU for MOI 5.

### Lentiviral transduction of MEFs

For experiments combining lentivirus and retrovirus during reprogramming, lentivirus was added on the second day of retroviral infection. Mixtures containing both viruses were made so that each well received 11 μL of each retrovirus, the calculated volume of each lentivirus for the desired MOI (based on functional titer), and DMEM + 10% FBS to reach a volume of 100 μL per 96-well. 5 μg/mL of polybrene was included to increase transduction efficiency. After adding the virus mixtures, the cells were spinfected by centrifuging the plate at 1500 × g for 30 minutes at 32°C. Reprogramming or twin plate protocols were then followed as described.

### Twin reprogramming assay

MEFs from the same frozen batch, and therefore biological source, were seeded onto two 96-well plates. Cells were transduced with retrovirus for two days, and with lentivirus on the second day of retroviral infection, as described here. The total lentiviral MOI was 20 across both viruses. For example, in conditions with mRuby2-RAS at MOI 5, the TagBFP MOI was 15. One biological twin was dissociated with trypsin at 4 dpi and flowed to measure mRuby2-RAS expression levels. The other biological twin was dissociated with DNase/papain at 14 dpi to measure iMN yield and purity.

### Reprogramming with signaling pathway inhibitors and PMA

The MEK inhibitor PD0325901 (Sigma-Aldrich, PZ0162), AKT inhibitor MK-2206 (Ambeed, A145592), JNK inhibitor SP600125 (Ambeed, A155219), and PMA (Ambeed, A175370) were dissolved in DMSO and stored at −20°C or −80°C. Before use, stock solutions were diluted in DMSO to reach 500x or 1000x the desired working concentration. This ensures equal DMSO concentrations across conditions. Each dilution was then added to the culture media immediately prior to use. All experiments with these small molecules began treatment at 1 dpi and continued through the experimental endpoint, with media changes every 2-3 days. For PMA experiments, RepSox was also added to all conditions beginning at 3 dpi. For immunofluorescent staining and western blot lysate, cells were treated with fresh media containing the inhibitors or PMA 20 minutes prior to fixation or lysis.

### CellTrace proliferation assay

At 1 dpi, cells were labeled with CellTrace™ Violet (CTV) dye from a CellTrace™ proliferation kit (Thermo Scientific, C34557). First, the CTV stock solution is diluted in PBS to a final concentration of 5 μM. After washing cells with PBS, 40 μL of the CTV working solution was added per 96-well, and the plate was incubated for 30 minutes at 37°C. After incubation, the CTV solution was removed and replaced with fresh DMEM + 10% FBS. Next cells were returned to the incubator or treated according to the experiment. At 4 dpi, cells were dissociated using trypsin and analyzed via flow cytometry. The hyperproliferative gate for each replicate was set based on the 20% of cells with lowest CTV signal in a control Puro infected condition.

### Flow cytometry

All flow experiments were performed with an Attune™ NxT flow cytometer. Cells were gated for live and single cells from the FSC and SSC channels using FlowJo™. Single cells were exported as csv files and analyzed using Python. A 405 nm laser with 440/50 filter was used for TagBFP and CTV. A 488 nm laser with a 510/10 filter was used for Hb9::GFP. A 561 nm laser with a 615/25 filter was used for mRuby2.

### Fixation and immunofluorescent staining

Cells were fixed with 4% paraformaldehyde for 1 hour at 4°C, then washed three times with PBS. Then cells were permeabilized with 0.5% Tween-20 in PBS for 1 hour at 4°C, followed by blocking in 5% FBS and 0.1% Tween-20 in PBS (blocking buffer) for 1 hour at 4°C. Next, cells were incubated with primary antibodies diluted in blocking buffer overnight at 4°C. The next day, cells were wash three times with 0.1% Tween-20 in PBS and incubated with secondary antibodies diluted in blocking buffer for 1 hour at 4°C. After three additional washes, nuclei were stained with 0.1 μg/mL DAPI in PBS for 30 minutes at room temperature. Finally, the DAPI solution was removed and cells were kept in PBS for imaging. Fluorescent images were taken using a Keyence™ All-in-one fluorescence microscope BZ-X800.

The primary antibodies used for immunofluorescent staining were: H-Ras Antibody (259) Alexa Fluor® 647 (1:50, Santa Cruz Biotechnology, sc-35, RRID:AB_627749); Phospho-p44/42 MAPK (Erk1/2) (Thr202/Tyr204) (1:250, Cell Signaling Technology, #9101, RRID:AB_331646); Phospho-Akt (Ser473) (1:50, Cell Signaling Technology, #9271, RRID:AB_329825). The secondary antibodies used here were: Donkey anti-Rabbit IgG (H+L) Alexa Fluor™ 647 for ppERK (1:250k, Thermo Fisher Scientific, A-31573, RRID:AB_2536183); Donkey anti-Rabbit IgG (H+L) Alexa Fluor™ 488 for pAKT (1:50k, Thermo Fisher Scientific, A-21206, RRID:AB_2535792).

### Senescence-associated β-galactosidase staining

β-gal staining was performed using a β-Galactosidase Staining Kit (Cell Signaling Technology, #9860) according to manufacturer’s instructions. In brief, cells were washed once with PBS, fixed with Fixative Solution for 15 minutes at room temperature, and washed three times with PBS. Next, 50 μL of 1x β-Galactosidase Staining Solution (prepared from 10x staining solution, 100x solution A, 100x solution B, and X-gal dissolved in DMSO, adjusted to pH 6.0) was added to each well. Plates were sealed with parafilm and incubated overnight in a dry incubator at 37°C. The next day, the staining solution was removed and replaced with PBS for imaging. Brightfield images of β-gal staining were captured using a Levenhuk M500 BASE camera attached to a light microscope.

### Western Blot

MEFs were seeded and infected as described here, except at a 6-well scale with amounts scaled up by well surface area. MEFs were seeded at 300k per 6-well, and 330 μL of each retrovirus was used per well. Cell lysate was collected at 4 dpi. To prepare lysate, cells were first washed with ice cold PBS, followed by the addition of 67 μL 1x RIPA buffer (Cell Signaling Technology, #9806) containing 1 mM PMSF (Cell Signaling Technology, #8553) per 6-well. Plates were incubated on ice for 5 minutes, then cells were detached using a cell scraper. Cells were sheared using blunt 21-gauge needles, and the lysate was clarified by centrifugation at 14000 × g at 4°C. Protein concentration was determined using a Bradford assay (Genesee Scientific, 18-442). Samples were separated using electrophoresis in a hand poured 12.5% bis-tris gel or a 4–15% Mini-PROTEAN® TGX™ precast gel (Bio-Rad, # 4561086), with 15 μg of total cell protein loaded per well. Proteins were transferred to a PVDF membrane using the iBlot™ 2 Dry Blotting System. Membranes were blocked with blocking buffer (5% milk and 0.1% Tween-20 in PBS) for 1 hour at room temperature with agitation. Then, membranes were incubated overnight with primary antibodies diluted in 10% blocking buffer. The next day, the membranes were washed with 0.1% Tween-20 in PBS followed by incubation with HRP-conjugated secondary antibodies in 10% blocking buffer for 1 hour at room temperature. After washing. HRP signals were detected using SuperSignal™ West Femto Maximum Sensitivity Substrate (Thermo Scientific, 34096), and blots were imaged using the ChemiDoc MP Imaging System. Western blot band intensities were quantified using the ImageJ gel analyzer tool.

The primary antibodies used in this study were: mouse anti-β-Actin (8H10D10) (1:50k, Cell Signaling Technology, #3700, AB_2242334); rabbit anti-Ras (G12V Mutant Specific) (D2H12) (1:2k, Cell Signaling Technology, #14412, RRID:AB_2714031); rabbit anti-Pan RAS (1:20k, Cell Signaling Technology, #3965, RRID:AB_2180216); rabbit anti-phospho-p44/42 MAPK (Erk1/2) (Thr202/Tyr204) (1:30k, Cell Signaling Technology, #9101, RRID:AB_331646); rabbit anti-p53 (D2H9O) (1:20k, Cell Signaling Technology, #32532, RRID:AB_2757821). The secondary antibodies used in the study were: goat anti-mouse IgG H&L (HRP) (1:50k, Abcam, ab205719, RRID:AB_2755049); goat anti-rabbit IgG H&L (HRP) (1:50k, Abcam, ab6721, RRID:AB_955447).

