## Supplementary Figures for "Chemogenetic tuning reveals optimal MAPK signaling for cell-fate programming"

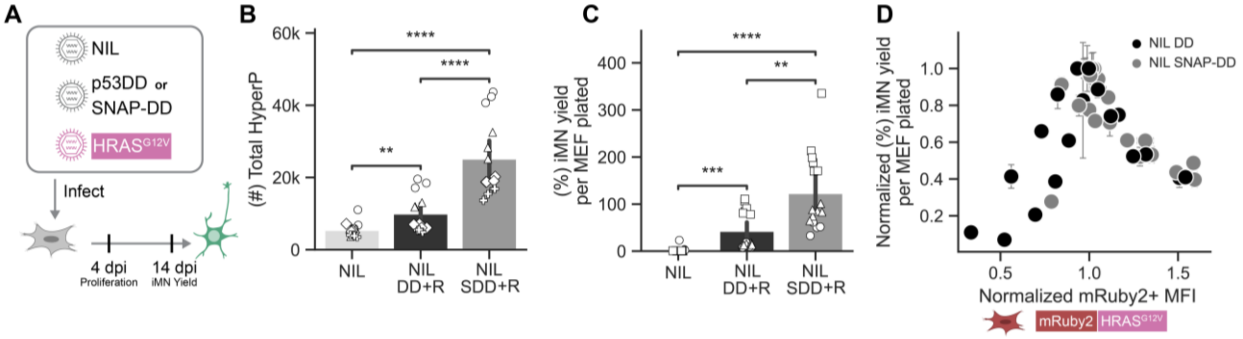


**Figure S1. Comparison of p53DD and SNAP-p53DD.**

A. Diagram showing experiments for comparing p53DD and SNAP-p53DD in conversion.

B-C. Total number hyperproliferative (HyperP) cells at 4 dpi and iMN yield at 14 dpi for NIL, NIL DD + HRAS$^{G12V}$ (R), or NIL SNAP-DD (SDD) + R. Mean is shown with 95% confidence interval; marker style denotes biological reps; n = 3 biological reps per condition.

D. Normalized iMN yield at 14 dpi vs. mRuby2-RAS+ geometric mean fluorescent intensity (MFI) at 4 dpi dpi for conditions with p53DD and SNAP-p53DD. Each point represents the mean of n = 3 technical replicates per bioreplicate ± standard error of mean (SEM). Marker color denotes p53DD (black) or SNAP-p53DD (grey); iMN yield is normalized so the maximum yield for each replicate overlays at 1.0 on the y-axis, and the mRuby2 levels are normalized so the mRuby2+ MFI at the MOI corresponding to the peak iMN yield overlays at 1.0 on the x-axis. n = 4 biological reps per condition

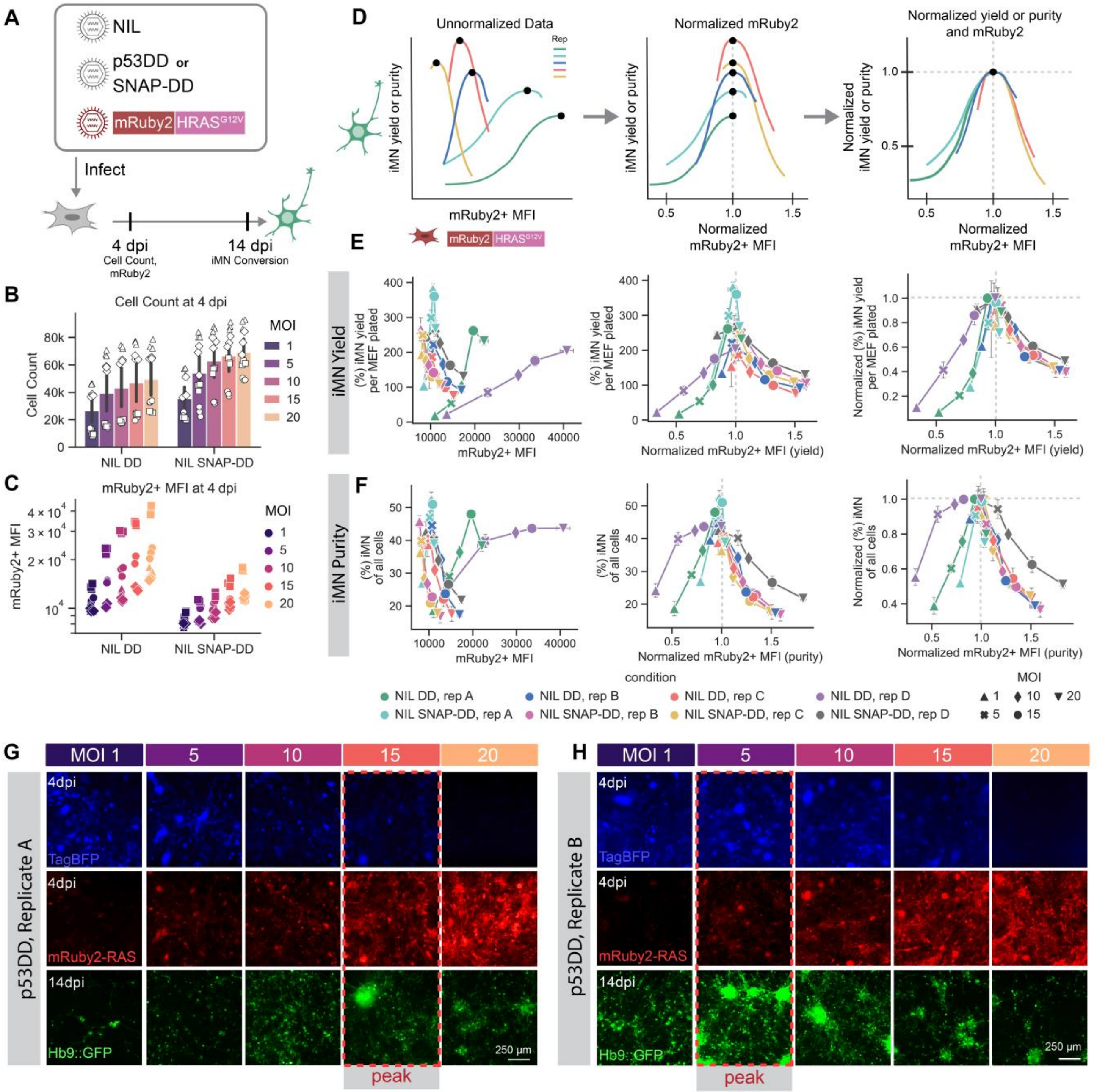


**Figure S2. Cell-fate programming responds biphasically to titration of HRAS$^{G12V}$**

A. Diagram showing experimental conditions for mRuby2-HRAS$^{G12V}$ titration experiments.

B-C. Cell count (B) and mRuby2+ MFI (C) at 4 dpi with mRuby2-HRAS$^{G12V}$ MOI titration in DD and SNAP-DD conditions. Marker style denotes biological replicates; n = 4 biological reps per condition.

D. Schematic depicting normalization strategy to correct for variance in mRuby2+ geometric mean fluorescent intensity (MFI) and conversion efficiency across p53DD conditions and biological replicates. First, for each DD condition and replicate, mRuby2+ MFI values are normalized to the MOI value that results in the peak iMN yield or purity. Next, the yield and purity is normalized by the maximum value for each DD condition and replicate. After normalization, the peak of each condition should align at x = y = 1.0.

E-F. iMN yield (E) and purity (F) at 14 dpi vs. mRuby2-HRAS$^{G12V}$ MFI at 4 days post infection (dpi) for conditions with p53DD and SNAP-p53DD at each stage of the normalization process. Each point represents the mean of n = 3 technical replicates ± standard error of mean (SEM). Marker style denotes MOI, hue denotes each combination of p53DD and biological replicate; n = 4 biological reps per condition.

G-H. Representative microscope images showing TagBFP and mRuby2-HRAS$^{G12V}$ expression at 4 dpi, and corresponding Hb9::GFP expression in iMNs at 14 dpi for two separate biological replicates. The mRuby2-HRAS$^{G12V}$ expression that correlates with peak iMN yield is indicated for each replicate. Scale bar represents 250 µm.

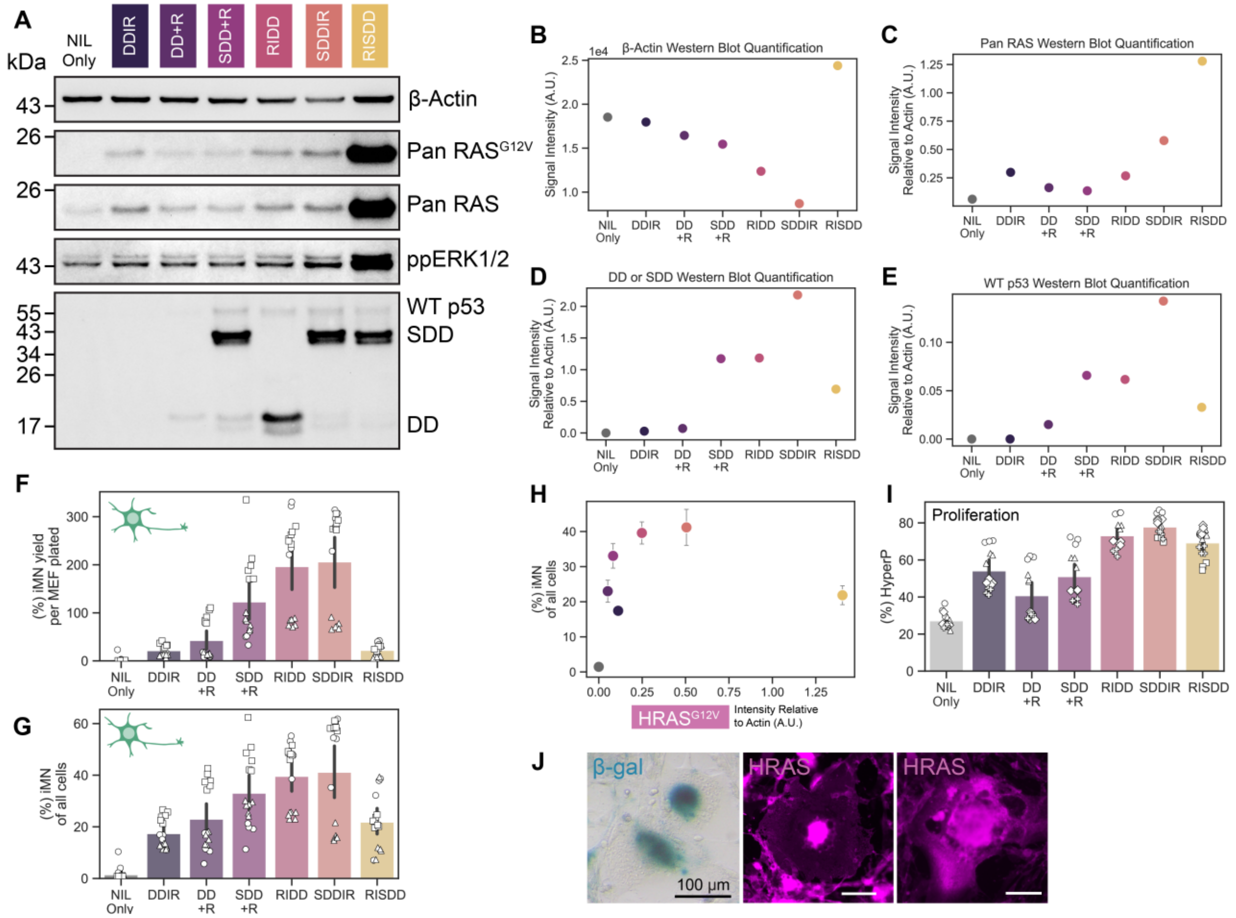


**Figure S3. HRAS$^{G12V}$ produces biphasic conversion through MAPK signaling and proliferation.**

A. Western blot of Pan RAS$^{G12V}$, Pan RAS, ppERK, wild type (WT) p53, SNAP-p53DD, and p53DD from lysate collected at 4 dpi with β-Actin as a loading control. All conditions include NIL. The p53 antibody recognizes all three p53 variants (WT p53, p53DD, SNAP-p53DD).

B-E. Quantification of western blots for β-Actin (B), Pan RAS (C), p53DD or SNAP-p53DD (D), and WT p53 (E) with each protein of interest normalized to β-Actin.

F-G. iMN yield and purity quantified at 14 dpi. Mean is shown with 95% confidence interval; marker style denotes biological reps; n = 3 biological reps per condition.

H. iMN purity at 14 dpi vs. Pan RAS$^{G12V}$ expression normalized to β-Actin levels measured from a western blot. Mean iMN purity is shown ± standard error of mean (SEM).

I. Percent HyperP at 4 dpi across conditions. Mean is shown with 95% confidence interval; marker style denotes biological reps; n = 4 biological reps per condition.

J. Microscope images showing examples of senescence morphology in NIL RISDD cells stained for senescence-associated β-galactosidase (β-gal) at 2 dpi and HRAS at 4 dpi. Scale bar represents 100 μm.

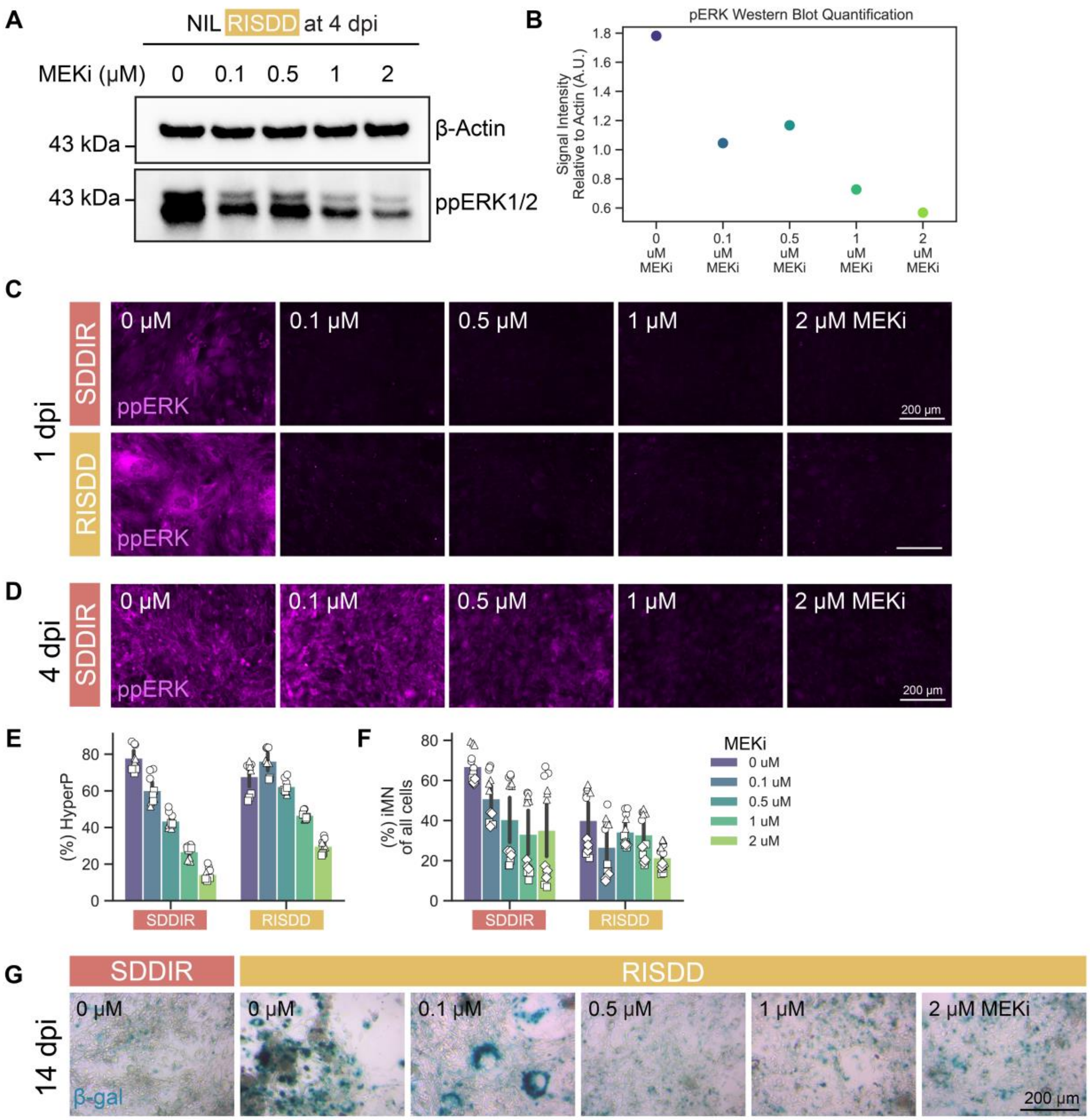


**Figure S4. Tuning MAPK signaling attenuates senescence and increases conversion.**

A. Western blot of ppERK from lysate collected at 4 dpi for NIL RISDD 20 minutes after treatment with fresh MEK inhibitor (PD0325901), including β-Actin as a loading control. MEKi was added to the media beginning at 1 dpi.

B. Quantification of western blot for ppERK normalized to β-Actin.

C-D. Representative images of immunofluorescent staining for ppERK at 1 and 4 dpi in the two polycistronic cassette conditions with the highest $HRAS^{G12V}$ expression (SDDIR and RISDD) with MEKi titration. MEKi was added to the media beginning at 1 dpi. Scale bar represents 200 µm.

E-F. Percent HyperP at 4 dpi and iMN purity at 14 dpi for with a MEK inhibitor titration for the two polycistronic cassettes with highest $HRAS^{G12V}$ expression. Mean is shown with 95% confidence interval; marker style denotes biological reps; n =3 biological reps per condition.

G. Representative images of cells stained for senescence-associated β-galactosidase (β-gal) at 14 dpi with MEKi titration. MEKi was added to the media beginning at 1 dpi. Scale bar represents 200 µm.

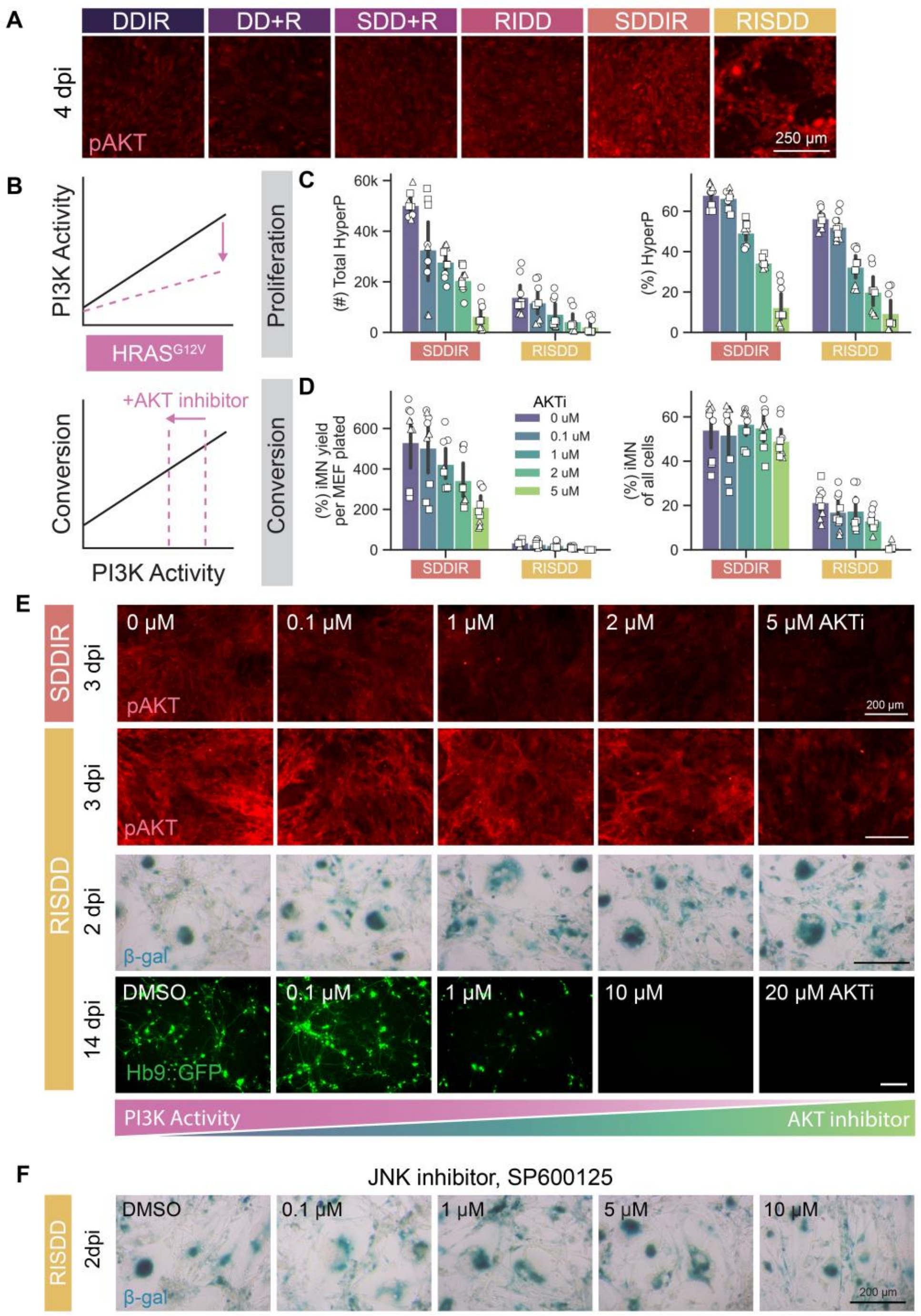


**Figure S5. Tuning of PI3K signaling does not attenuate senescence or increase the rate of conversion.**

A. Representative images of immunofluorescent staining for pAKT at 4 dpi across DD HRAS$^{G12V}$ polycistronic cassette conditions. Scale bar represents 250 μm.

B. Diagram depicting expected results of adding an AKT inhibitor on PI3K signaling levels and conversion.

C. Total number HyperP and percent HyperP at 4 dpi with an AKT inhibitor (MK-2206) titration for the two polycistronic cassettes with highest HRAS$^{G12V}$ expression (SDDIR and RISDD). AKTi was added to the media beginning at 1 dpi. Mean is shown with 95% confidence interval; marker style denotes biological reps; n = 3 biological reps per condition.

D. Conversion yield and purity quantified at 14 dpi with an AKT inhibitor titration for the two polycistronic cassettes with highest HRAS$^{G12V}$ expression. AKTi was added to the media beginning at 1 dpi. Mean is shown with 95% confidence interval; marker style denotes biological reps; n = 3 biological reps per condition.

E. Representative images of cells stained for pAKT at 3 dpi in SDDIR and RISDD polycistronic cassettes, and senescence-associated β -galactosidase (β-gal) at 2 dpi and Hb9::GFP expression in iMNs at 14 dpi for the polycistronic cassette with the highest HRAS$^{G12V}$ expression (RISDD) with AKTi titration. Scale bar represents 200 μm.

F. Representative images of cells stained for senescence associated β-gal at 2 dpi in RISDD condition after treatment with JNK inhibitor (SP600125) beginning at 1 dpi. Scale bar represents 200 μm.

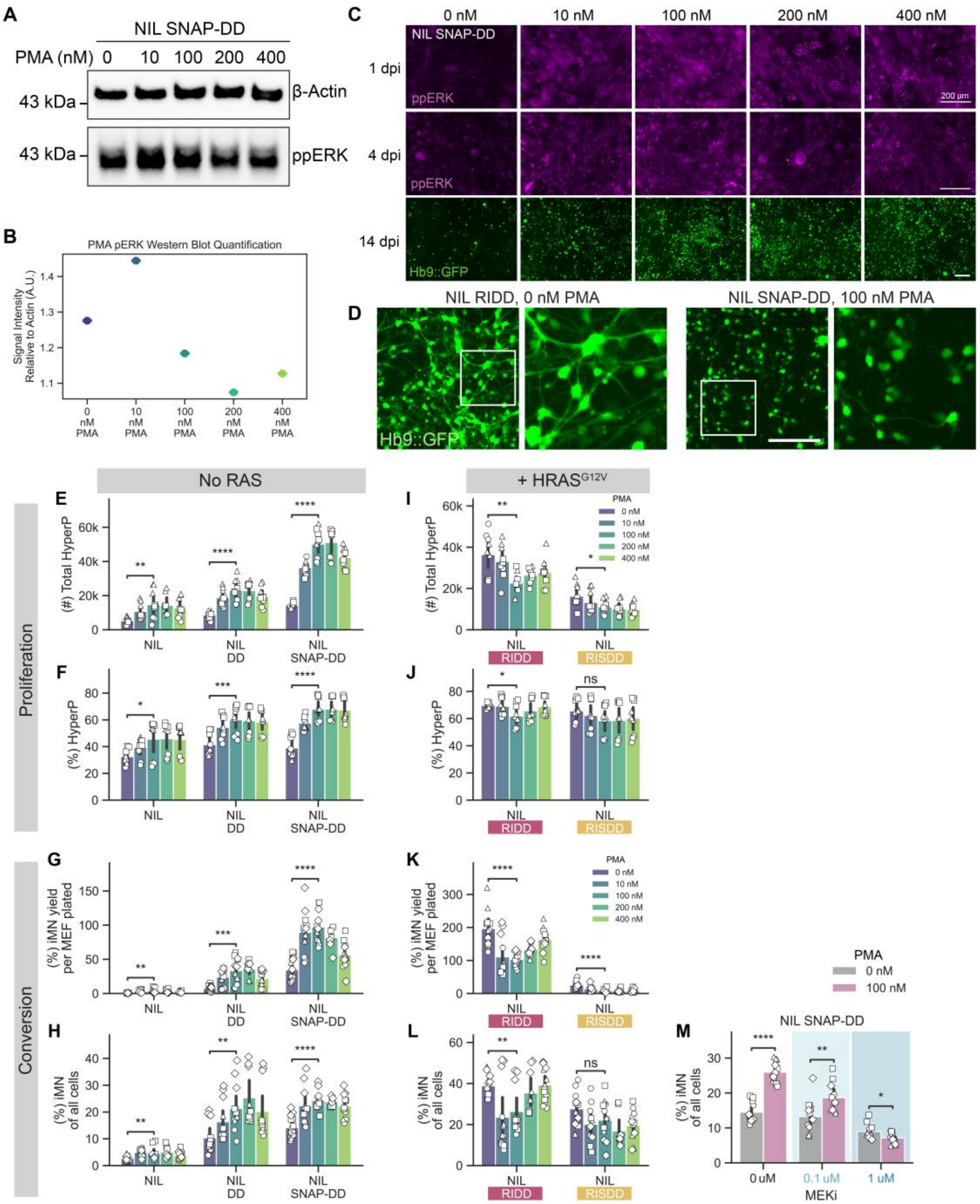


**Figure S6. Activation of MAPK signaling induces high rates of conversion in the absence of mutant RAS.**

A. Western blot of ppERK from lysate collected at 4 dpi for NIL SNAP-p53DD 20 minutes after a media change with fresh PMA including β-Actin as a loading control.

B. Quantification of western blot for ppERK normalized to β-Actin.

C. Representative images of immunofluorescent staining for ppERK in NIL SNAP-DD infected cells at 1 dpi or 4dpi that were fixed 20 minutes after replacing media with fresh PMA at varying concentrations, and Hb9::GFP+ iMNs at 14 dpi. Scale bar represents 200 µm.

D. Representative images comparing iMN morphology at 14 dpi for NIL RIDD without PMA to NIL SNAP-DD + 100 nM PMA. Scale bar represents 200 µm.

E-H. Total number and percent HyperP cells at 4 dpi with a PMA titration for conditions with and without HRAS$^{G12V}$. Mean is shown with 95% confidence interval; marker style denotes biological reps; n = 3 biological reps per condition; t-test independent samples.

I-L. Conversion yield and purity at 14 dpi with a PMA titration for conditions with and without HRAS$^{G12V}$. Mean is shown with 95% confidence interval; marker style denotes biological reps; n = 4 biological reps per condition; t-test independent samples.

M. Conversion purity quantified at 14 dpi for cells infected with NIL SNAP-DD and treated with combinations of 0 nM or 100 nM PMA to activate MAPK signaling and 0 µM, 0.1 µM, or 1 µM of MEK inhibitor (PD0325901) to inhibit MAPK signaling. Small molecules were added to the media beginning at 1 dpi. Mean is shown with 95% confidence interval; marker style denotes biological reps; n = 4 biological reps per condition; t-test independent samples.

Significance summary: p > 0.05 (ns); *p ≤ 0.05; **p ≤ 0.01; ***p ≤ 0.001; and ****p ≤ 0.0001.
